# Repeated head-exposures to a 5G-3.5 GHz signal do not alter behavior but modify intracerebral gene expression in adult male mice

**DOI:** 10.1101/2024.12.13.628345

**Authors:** Julie Lameth, Juliette Royer, Alexandra Martin, Corentine Marie, Délia Arnaud- Cormos, Philippe Lévêque, Roseline Poirier, Jean-Marc Edeline, Michel Mallat

**Affiliations:** Institut du Cerveau, ICM, Inserm U 1127, CNRS UMR 7225, Sorbonne Université, F-75013, Paris, France; Université Paris-Saclay, CNRS, Institut des Neurosciences Paris-Saclay, 91400, Saclay, France; Univ. Limoges, CNRS, XLIM, UMR 7252, 123 avenue Albert Thomas, F-87000 Limoges, France; Institut Universitaire de France (IUF), 1 rue Descartes, 75005 Paris, France

**Keywords:** 5G, electromagnetic field, memory, behavior, transcriptome, mitochondria, glutamatergic synapse

## Abstract

The 5^th^ generation (5G) of mobile communications promotes human exposures to electromagnetic fields exploiting the 3.5 GHz frequency band. We have analyzed behaviors, cognitive functions and gene expression in mice submitted to asymmetrical head-exposures to a 5G-modulated 3.5 GHz signal. The exposures were applied 1h daily, 5 days per week over a six-week period, at a specific absorption rate (SAR) averaging 0.19 W/kg over the brain. Locomotor activity in an open-field, object-place and object recognition memories were assessed repeatedly after four weeks of exposure and did not reveal any significant effect on the locomotion/exploration, anxiety level or memory processes. mRNA profiling was performed at the end of the exposure period in two symmetrical areas of the right and left cerebral cortex in which the SAR values were 0.43 and 0.14 W/kg, respectively. We found significant changes in the expression of less than 1% of the expressed genes with over-representations of genes related to glutamatergic synapses. The right cortical area differed from the left one by an over-representation of responsive genes encoded by the mitochondrial genome. Our data show that repeated head-exposures to a 5G-3.5 GHz signal can trigger mild transcriptome alterations without change in memory capacities or emotional state.

## Introduction

Wireless communications generate a worldwide exposure of people to radiofrequency (**RF)** electromagnetic fields (**EMF**), which have raised concerns and debates on the public health impact of these physical agents. The common use of mobile phones promotes head exposures of users, which have stimulated a wealth of investigation aiming at determining possible effect of EMF, implemented by second, third and fourth generations (G) of mobile or wireless (WIFI based on IEEE 802.11) communications, on the central nervous system (CNS). The influence on behavioral or cognitive functions were investigated on human subjects and animal models [1–3].

Among these, rodent models allow assessment of spontaneous behaviors and emotional states or cognitive capacities such as learning and memory and allow analyzing the cellular and molecular mechanisms responsible for behavioral modifications induced by environmental agents. Regarding the effects of RF-EMF on the learning abilities and spatial memory of rodents, published works show heterogeneous results, reporting either no effect, or in contrast, significant modifications of learning or memory abilities [3–7]. The diversity most likely reflects the influence of a combination of experimental parameters that vary across studies. Parameters related to exposure conditions include frequency and modulation of the signals, exposure mode limited to the head or extended to the whole body, its duration and periodicity, as well as the energy absorbed at the level of the brain or the whole body, which is quantified by the specific absorption rate (**SAR**) expressed in W/kg.

Although 2G, 3G and 4G systems remain on duty, the use of 5G mobile communications is expanding fast and was expected to reach 2.3 billions subscriptions by the end of 2024 (Ericson mobility report, 2024; https://www.ericsson.com/en/reports-and-papers/mobility-report). 5G telecommunications systems exploit RF bands not used in previous systems around 3.5 GHz or 26 GHz. In contrast with exposures to millimetric 26 GHz RF that are fully absorbed within 1-2 mm thick skin or underneath tissues, a head exposure to 3.5 GHz RF results in energy transfer that can reach brain cells [8,9]. So far, the effect of 3.5 GHz RF exposure on the CNS remains little investigated. A recent electroencephalographic assessment of healthy human adults submitted to 17 min far-field exposure to a 5G-3.5 GHz signal revealed no significant overall changes in the electrical brain activity, when the brain-averaged SAR was estimated to a value as low as 0.008 mW/kg. However, alterations in power spectral densities occurred at the level of some electrodes during the exposure or within a 17 min time period following exposure, which depended on eye open or closed conditions [8]. No clear changes in the spontaneous electrophysiological activity of neuronal networks could be observed when using primary cultures of embryonic mouse neurons that were exposed to a 5G-3.5GHz signal applied for 15 min at SAR levels of 1 W/kg or 3 W/kg [10]. Besides studies focused on neuronal activities, the behavioral and molecular consequences of 3.5 GHz RF were investigated in different animal models. Exposures of adult male guinea pigs applied for three days at whole body SAR values of 2, 4 or 10 W/kg, did not alter locomotor activities or hearing thresholds but triggered molecular and ultrastructural marks of oxidative stress at the level of the auditory cortex [11]. Short-term exposures with SAR value estimations ranging from 0.0026 to 0.26 W/Kg were found to increase activity and reduce sleep duration in male flies, whereas parental and embryonic exposures had opposite behavioral effects in F1 male offsprings. Behavioral changes were associated with altered expression of genes encoding heat shock proteins or circadian clock components and modifications in the levels of excitatory and inhibitory neurotransmitters [12]. Parental and embryonic exposures of flies also resulted in shortening in the mean development time, together with increased activity of antioxidant enzymes and modifications of the fly microbiota assessed at late larval stages [13]. Developmental impacts were also documented in the Zebrafish, showing that exposures during embryonic life can result in hypoactivity or anxiety related-behaviors at larval or adult stages [14–16]. Transcriptome profiling of zebrafish embryos performed at the end of the RF exposure revealed significant changes in transcript levels limited to 28 genes involved in metabolic pathways [15].

In the current study, we have set up an experimental mouse model to evaluate whether a one-month chronic exposure to 5G-3.5 GHz signal (1 hour / dayf 5 days a week), at adulthood, can alter exploration, anxiety, memory and intra-cerebral gene expressions. The exposures were performed in awake, head-restrained, animals to ensure a daily head exposure at a power set to limit upper SAR values around 0.5 W/kg within the CNS. After 5G-exposure, the locomotor activities in open-field (**OF**) and learning and memory abilities of mice tested in novel object recognition and object location tasks were compared to those of pseudo-exposed mice, which were also awake, head-restrained, at the same time than the exposed mice but the 5G-3.5 GHz signal was turned off.

Unrestrained cage control mice were also used to evaluate the potential deficit induced by the restrained conditions. After completion of the behavioral task, the brains were quickly removed from the skull and a whole genome mRNA profiling was performed in symmetrical areas of the right and left cerebral cortex, which differed owing to the local energy deposit as indicated by our dosimetric analysis.

## Methods

The experiments were performed under the national license A-91-557 (APAFIS project # 33606 from the “Direction générale de la recherche et de l’innovation”) using procedure No. 32-2011 validated by the Ethics Committee in Animal Experimentation (CEEA 59, Paris Centre et Sud). All procedures were performed in accordance with the guidelines established by the European Communities Council Directive (2010/63/EU Council Directive Decree).

### Subjects

Seven weeks old C57BL/6J male mice (n=32) were obtained from Charles River (L’Arbresle, France). They were housed in a humidity (50–55%) and temperature (22– 24°C)-controlled animal facility on a 12 h/12 h light/dark cycle (light on at 7:30 A.M.) with free access to food and water. After one week of familiarization with the animal facility, two groups of mice underwent a brief (30min) surgery under isoflurane anesthesia (1.5- 2%) to secure a small plastic hook (0.25g) to the skull bone with super-bond C&B (Sun Medical) and dental acrylic cement (Methax, Makeval LTD). Metacam (0.1mg/kg, i.p.) was administrated after the surgery to help the animal recovery. This small hook allowed to maintain the animals in restrained conditions during the exposure. Each mouse was daily weighted and all mice included in this study gained weight after this initial surgery. A third group of control (**CTRL**) unrestrained mice was used to evaluate the potential behavioral alteration induced by the restrained conditions.

### Exposure system and exposure protocol to 5G-3.5 GHz signals

Head-only exposures to 5G-3.5 GHz signal were performed in awake restrained mice. Each mouse was habituated to the exposure conditions by progressively increasing the time during which the mouse was head-fixed (from 15 min to 60 min over a week) in a red plastic tube (internal diameter: 32 mm) with the hook screwed to a small plastic post (see Figure 1A-B). Each day, a dipole antenna (SID3500, MVG, France) was positioned 5 mm from the animal head in a fixed/standardized position (Figure 1B) during one hour at vicinity of the right temporal cortex.

**Figure 1.**
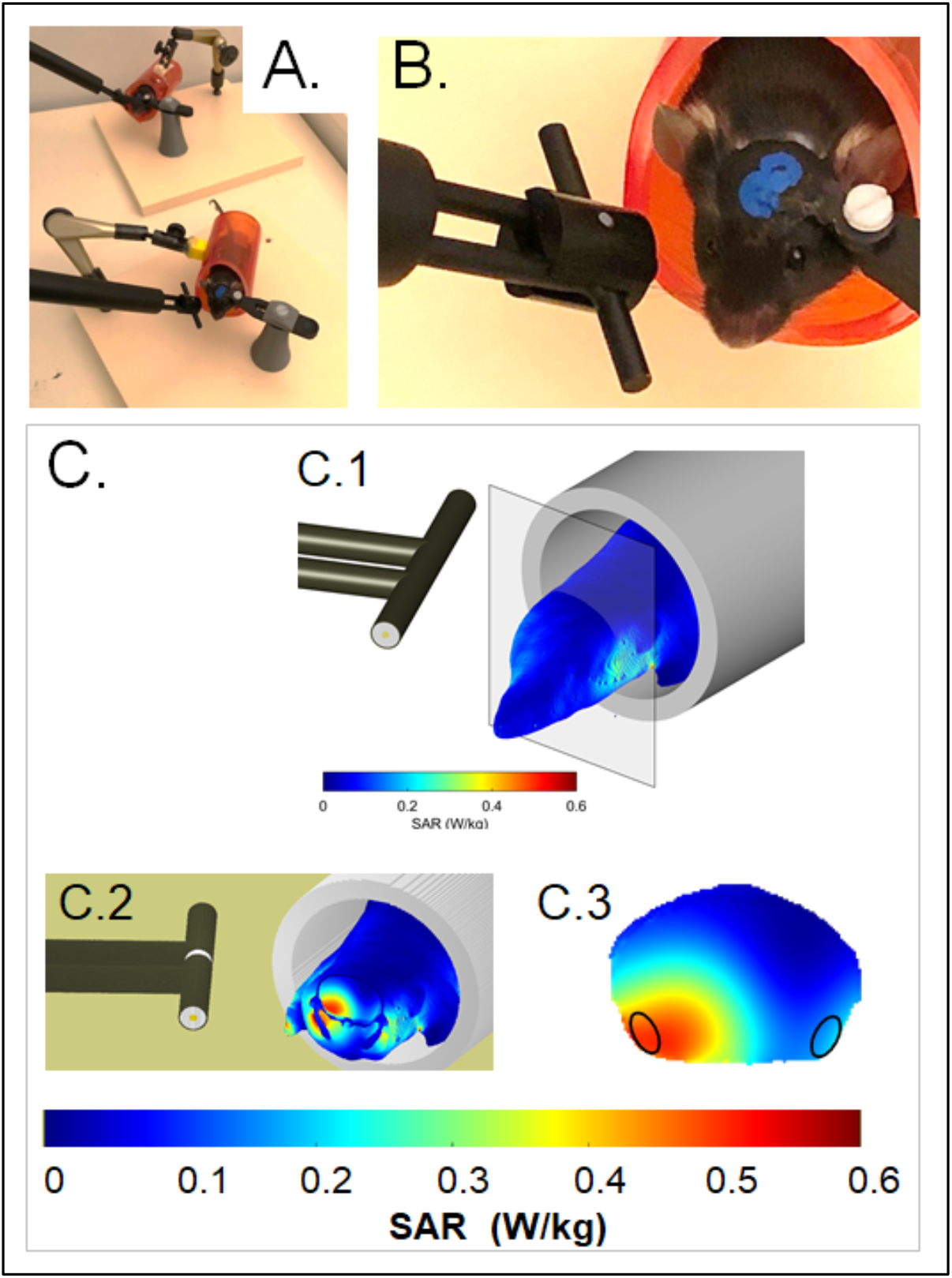
exposure setup and dosimetry. (A,B) Head restrained mice in red plastic tubes with dipole antennas positioned at vicinity of the right temporal cortex. The low magnification view (A) shows two matched 5G-exposed and pseudo-exposed animals (the antenna of the pseudo -exposed mouse is not connected to the 5G-3.5 GHz generator). (C) Dosimetric analysis of specific absorption rates (SAR) in the mouse head and trunk. A heterogenous model of mouse and dipole antenna was used to evaluate the local SAR in the head and trunk at a spatial resolution of 0.2 mm^3^. (C1) 3D view illustrating SAR values at the surface of the head and trunk. (C2) 3D view with a coronal section showing external and internal distributions of the SAR values in the head. The area of the sectioned brain is delimited by a black contour. (C3) 2D view focused on the coronal brain sections. Areas of the right and left entorhinal-piriform cortex, which were sampled for transcriptomic analyses, are encircled. Bars shows the color-code scale of SAR values

The exposure system was similar to the one described in a previous study [17], replacing the frequency emitted by the radiofrequency generator to generate a 5G-3.5 GHz signal corresponding to the 5G NR (release 15, Digital Standards SMBVB-K444; Rohde & Schwarz) with FDD duplexing, QPSK modulation and 100 MHz channel bandwidth. Briefly, a radiofrequency generator emitting a 5G-3.5 GHz electromagnetic field (SMBV100B, Rohde & Schwarz, Germany) was connected to a power amplifier (ZHL-4W-422+, Mini-Circuits, USA), a circulator (Pasternack, PE83CR1005, CA, USA), a bidirectional coupler (Mini-circuits, ZGBDC30-372HP+, NY, USA) and a four ways power divider (Mini-circuits ZB4PD-462W-N+, NY, USA), allowing potential simultaneous exposure of four animals. A powermeter (E4417A and E9323A, EPM-P Series Power Meter, Agilent, USA) connected to the bidirectional coupler allowed continuous measurements and monitoring of incident and reflected powers within the setup.

Each exposed mouse was matched with a pseudo (**PSD**)-exposed mouse that was in head-fixed restrained conditions next to it (at about 30 cm, Figure 1A). An antenna was also placed at 5 mm from the head of the PSD-exposed mice but this antenna was not connected to the 5G-3.5 GHz generator.

### Dosimetry

Similar to previous studies [18–20], specific absorption rates (SARs) were determined numerically using a numerical mouse model with the Finite Difference Time Domain (**FDTD**) method [21–23] with a spatial resolution of 0.2 mm^3^. SAR was also determined experimentally in an homogeneous mouse model using a Luxtron probe for measurement of rises in temperature. In this case, SARs, expressed in W/kg, were calculated using the following equation: SAR = CΔT/Δt with C being the calorific capacity in J/(kg.K.), ΔT, the temperature change in °K and Δt, the time in seconds. Numerically determined SAR values were compared with experimental SAR values obtained using homogenous models, especially in the equivalent mouse brain area. The difference between the numerical SAR determinations and the experimentally detected SAR values was less than 30% as recommended by standard EMF exposure guidelines.

Figure 1C shows the SAR distribution in the mouse head and anterior trunk with a mouse model, which matched that of the mice in terms of weight and size. Intra cerebral SAR values (Figure 1C.3) were the highest in ventral cortical areas ipsilateral to the antenna, which covered adjacent area of the right entorhinal and piriform cortex (Ent-Pir Cx), reaching values of 0.43 ± 0.12 W/kg (volume-averaged value ± SD among the SAR voxel values over the defined volume). SAR values in the contralateral (left) Ent-Pir Cx dropped to 0.14 ± 0.05 W/kg. The whole brain-averaged SAR was 0.19 ± 0.12 W/kg. As the weight of the exposed mice was homogenous, differences in tissues thickness at the level of the head were probably negligible, so the actual SAR in the cerebral cortex was very similar from one exposed animal to another.

### Behavioral protocol

Twenty days after the beginning of the 5G exposure (Figure 2A), we assessed spontaneous locomotion, exploratory behavior and potential anxiety in an OF arena, then long-term recognition memory using object recognition task. These behavioral tasks were performed 10 min after the exposure or pseudo-exposure session. The testing procedure was similar to the one used in [24]. The protocol started by three habituation sessions to explore an OF (Figure 2B). The square OF (50x50cm) had black sidewalls (35 cm high) and its floor was covered with 2 mm of sawdust. A camera connected to a video-tracking system (ANY-maze^TM^ software, Stoelting) was placed above the OF to record the mice activity. Experiments were undertaken under homogeneous dim illumination (<50 lux). On days 1-3, each mouse was allowed to explore the empty OF for 20 minutes (day 1) or 10 minutes (days 2 and 3). During these three sessions, the total travelled distance, the rearing number and percent distance traveled in the peripheral area (5 cm from the walls) were recorded to evaluate locomotion, exploration and also potential anxiety revealed by thigmotaxis (tendency of an anxious mouse to remain close to walls). To assess novelty-seeking behavior, on day 4, two identical rectangular plastic objects were placed in the center zone and the mice could freely explore them for 10 min. The time exploring the objects as well as the travelled distances and thigmotaxis were recorded. Object exploration was defined as the mouse touched the object with its nose, or sniffed it with a distance between the nose and the object less than 2 cm. The following days (days 5-8), mice were tested in two successive experiments: the novel object recognition task (**NOR**) to assess object recognition memory and the object location task (**OL**) to assess spatial memory (Figure 2B). The order of these experiments was counterbalanced among individuals and groups (control mice, exposed mice, pseudo-exposed mice). Each experiment consisted of acquisition session followed by a retention test 24 hours later (Figure 2B). Thus, on day 5 or 7, three different objects were placed in the OF and could be freely explored during three consecutive trials of 5 minutes with a 5-minute inter-trial interval (**ITI**). The objects were plastic toys of different colors and shapes (3–6 cm diameter, 3–6 cm high). The objects and their spatial arrangement in the test box were chosen in a pseudorandom order and were counterbalanced between mice. Twenty-four hours later (days 6 and 8), a memory test (one trial of 5 minutes) was performed (Figure 2B). For these tests, because mice are spontaneously attracted by novelty, either one object explored during previous acquisition session was replaced by a novel object to assess object recognition memory, either one of the 3 objects explored during the previous acquisition session was displaced to a novel location to assess object place memory (days 6 and 8, Figure 2B). For each session (acquisition and retention), the time spent exploring each object was analyzed, as well as the total object exploration and total travelled distance. For each subject, retention performance was expressed as the percent time spent exploring the novel /displaced object over the total object exploration time during the 5-minute-session.

**Figure 2.**
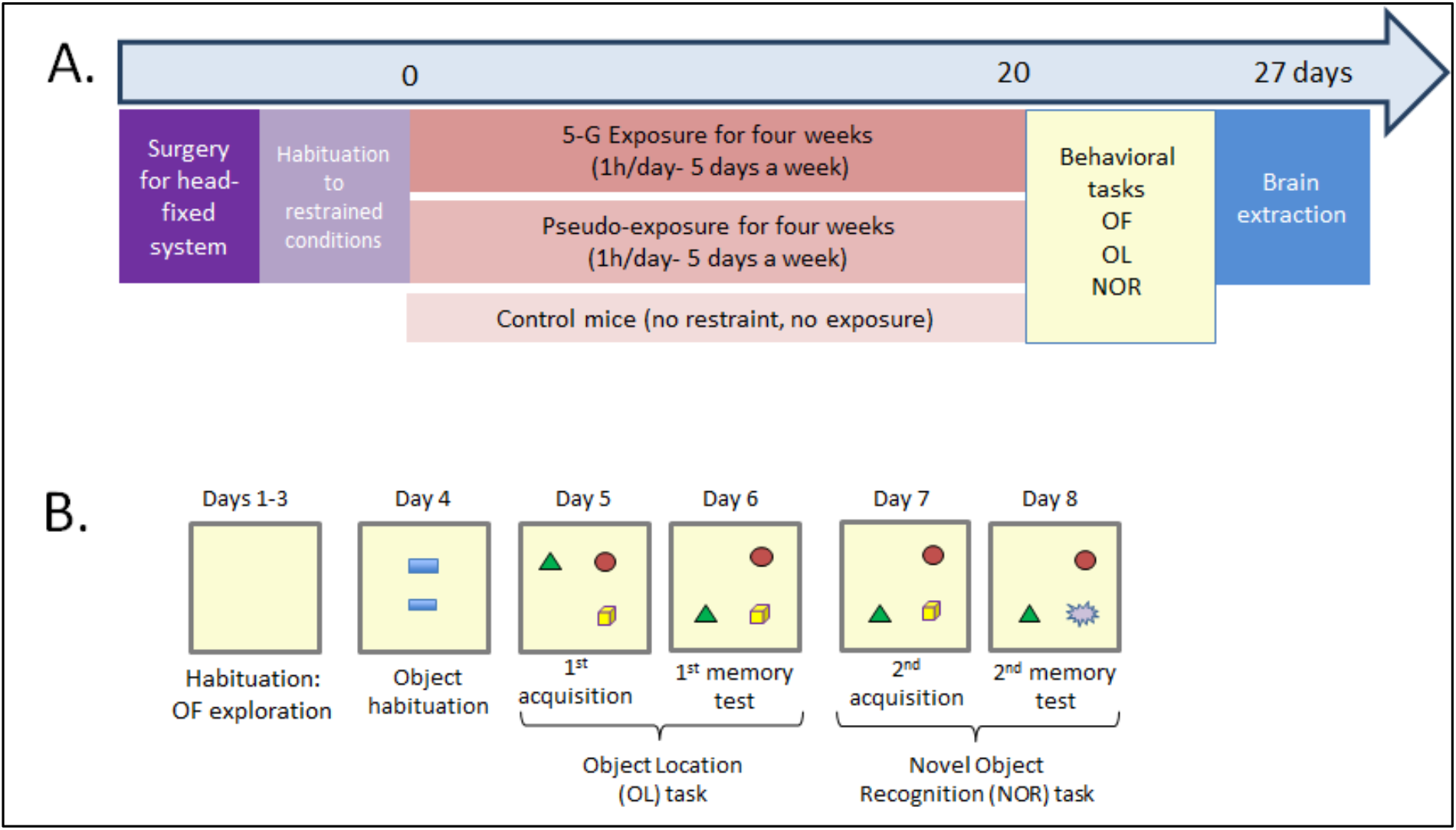
Timeline of the experimental procedures. (A) After surgery and habituation to restrained conditions, mice were exposed to 5G-3.5 GHz signal, 1 hour per day, 5 days a week, during 4 weeks. After day 20, mice were submitted to behavioral tasks after each 1h-exposure session. As controls, the behavior of a group of mice kept under restraint conditions during the same period (Pseudo exposed group) and a group of control mice without exposure or restraint was also assessed. At the end, brains of 5G- and Pseudo- exposed mice were collected for biochemical analysis. (B) Behavioral tasks used to assess exploration/locomotion in an open-field (OF) and Object recognition memory with Object Location (OL) and Novel Object Recognition (NOR) tasks. Mice were allowed to explore OF during 3 sessions (days 1-3). On day 4, they could explore 2 objects placed in the OF. From day 5, they were submitted to memory tasks. For OL task, one object has been displaced (here, the green triangle). For NOR task, an object has been replaced by a new one. In both conditions, because mice are spontaneously attracted by novelty, their recognition memory is assessed.

### Statistical analyses

For all behavioral statistical analyses, GraphPad Prism (version 8.0.2) was used. The data are presented as the group means ± SEM. For each parameter collected during the behavioral tests, the normality of the distribution and the homogeneity of variance were checked before running the statistical analyses (respectively, Shapiro-Wilk test and Levene’s test). Differences among groups was assessed by the one-way analysis of variance (**ANOVA**), or two-way ANOVA for multifactorial analyses. As appropriate, Bonferroni’s *post hoc* tests were employed to determine group differences. For memory test, one sample t-test was used to compare the exploration of the displaced/novel object to chance level (33.33%). The significance level was set at P<0.05.

### Tissue preparation for transcriptome analyses

Mice were killed by decapitation under isoflurane anesthesia 24h after the last behavioral testing. The brains were quickly removed from the skulls and frozen. Tissue samples were extracted from a brain region covering adjacent area of the entorhinal and the piriform cortex (**Ent-Pir Cx**) of the right and the left hemispheres in 100 µm-thick coronal sections cut on a cryostat and distributed over a cortical region starting 9 mm caudal to the anterior part of the olfactory bulb and extending over 1 mm in the rostro-caudal axis. Tissues were stored frozen at -80 °C until use.

### RNAseq and data analysis

Total RNA was extracted from the Ent-Pir Cx collected from the right and left hemispheres of PSD-exposed (n= 8) or 5G-exposed mice (n=7) using RNeasy® Micro kit (QIAGEN). Quality of RNA was confirmed on Agilent TapeStation (RIN > 8). RNA seq libraries were prepared using TruSeq® Stranded mRNA LT kit (Illumina) and sequenced with Illumina NovaSeq 6000 platform (double stranded 2 x 43 million reads with 2 x 100 nucleotides length per sample).

Quality of raw data was evaluated with FastQC [25]. Poor quality sequences and adapters were trimmed or removed with fastp tool, with default parameters, to retain only good quality paired reads [26]. Illumina DRAGEN bio-IT Plateform (v3.10.4) was used for mapping on mm10 reference genome and quantification with gencode vM25 annotation gtf file. Library orientation, library composition and coverage along transcripts were checked with Picard tools. Following analyses were conducted with R software. Data were normalized with DESeq2 (v1.34.0) bioconductor package [27], prior to differential analysis with glm framework likelihood ratio test from DESeq2 workflow. Multiple hypothesis adjusted p-values were calculated with the Benjamini-Hochberg procedure to control false discovery rate (**FDR**). Results were considered statistically significant for FDR- adjusted p-values ≤ 0.05 and the fold-change (FC) threshold was set to 1.2 to determine differentially expressed genes (**DEG**). Clusterings have been performed using “dist” and “hclust” functions in R, using Euclidean distance and Ward agglomeration method. Finally, over-representation enrichment analysis was conducted with DAVID Functional annotation Tool [28,29](Huang et al., 2009; Sherman et al., 2021). GO terms were considered as enriched if fold enrichment ≥ 2.0, FDR ≤ 0.05 and minimum number of regulated genes in pathway/term ≥ 2.0.

## Results

After 20 daily sessions of 1h-exposure, we assessed the effects of 5G-3.5 GHz signal on several behavioral parameters and memory processes for six additional days. During this period, the daily 5G/PSD-exposure sessions continued. Mice were behaviorally tested after 10 min of rest in their home/personal cage. Because head-fixed conditions could also induce behavioral alterations such as hyperactivity or increased anxiety, we also compared the performances of 5G and PSD-exposed mice with CTRL mice (without restraint) for these parameters. For behavioral studies, four 5G-exposed animals had to be retired from analysis because their plastic hook damaged during the last sessions: two before behavioral tasks, one before the NOR task and one before the OL task. Therefore, the results presented below are from 8 CTRL mice, 10/11 5G-exposed mice and 11 PSD- exposed mice.

### Chronic 5G-exposure does not lead to hyperactivity and abnormal anxiety

During the first session in OF, 5G-exposed mice showed normal locomotion and exploration activity and the total travelled distance was similar to those of PSD-exposed and CTRL mice (Figure 3A; F(2,28)= 1.64; p=0.21). Over the 20 minutes of test, the travelled distance progressively decreased, revealing normal habituation to this novel environment and no difference between groups was found (Figure 3B; 5-min-session effect: F(3,84)=134; P<0.0001; group effect: F(2,28)=1.64; P=0.21, 5-min session x group: F(6,84)=0,63; P=0.7). Similar results were observed during the 2^nd^ and 3^rd^ session of habituation in OF with no difference between groups (Supplementary Table S1). Regarding another index of exploratory behavior, our analyses revealed that the numbers of rearing of 5G- and PSD-exposed mice were significantly increased compared to CTRL mice during the first 20-min session in OF (Figure 3C; F(2,28)=12.72; P=0.0001; Bonferroni’s multiple comparisons test: CTRL vs 5G: P=0.0003; CTRL vs PSD: P=0.0004). No difference was found between 5G and PSD-exposed mice (P>0.99). This increase of number of rearings could be due to daily manipulation of the 5G-exposed and PSD- exposed mice in contrast to CTRL mice. This interpretation is reinforced by the data observed during the 2^nd^ and 3^rd^ sessions where the numbers of rearing were similar between the three groups (Supplementary Table S1). Finally, to assess whether 5G- exposure could induce anxiety-like behaviors, we analyzed the percentage of distance travelled in the central zone of the OF. For this parameter, the 5G- and PSD-exposed mice travelled similar distance in the central area of the OF. However, as shown in Figure 3D, CTRL mice travelled significantly less distance in this central area compared to exposed mice (Figure 3D for the 1^st^ session, F(2,28)=6.65; P=0.0043; Bonferroni’s multiple comparisons test: 5G *vs* PSD: P>0.99; CTRL *vs* 5G: P=0.0078; CTRL *vs* PSD: P=0.01). From the 2^nd^ session in OF, there was no difference for this parameter between the three groups (supplementary data for 2^nd^ and 3^rd^ session, Table S 1), confirming that, during the 1^st^ session in OF, CTRL mice were slightly more anxious than 5G- and PSD-exposed, daily manipulated, mice. We observed similar results when two objects were introduced into the OF for the first time. During this session (day 4), there was no difference in time spent to explore the objects between 5G- and PSD-exposed mice, but CTRL mice spent significantly less time exploring the two objects than exposed mice (Suppl. Figure 1A: F(2,27)=6.059; P=0.0067; Bonferroni’s multiple comparisons test: 5G *vs* PSD: P>0.99; CTRL *vs* 5G: P=0.01; CTRL *vs* PSD: P=0.017). Together, our analyses suggest that 5G- exposed mice did not display hyperactivity or abnormal anxiety, and their levels of exploration of OF/ objects were similar to the PSD-exposed mice.

**Figure 3.**
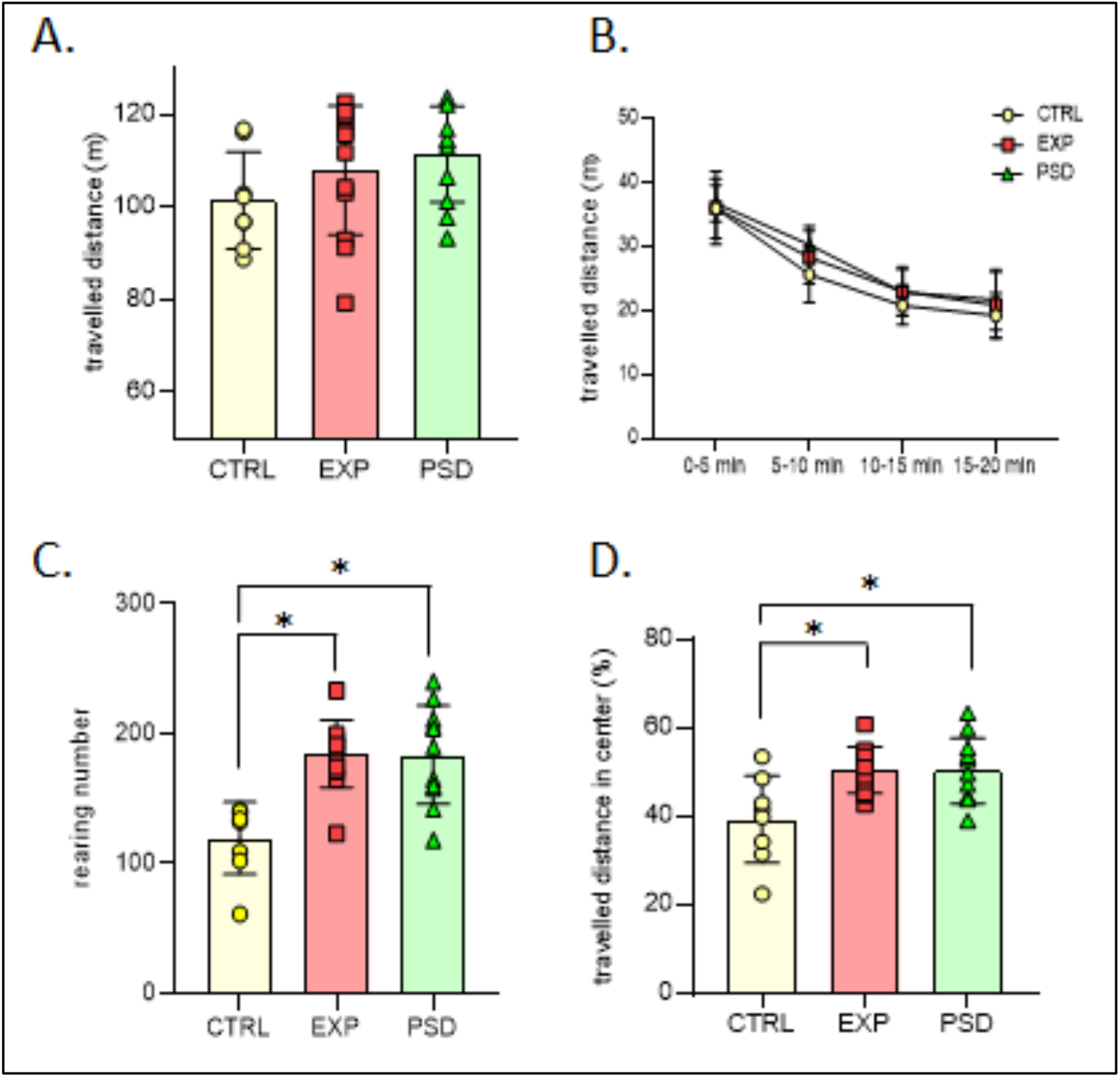
Behavioral parameters observed during the 1^st^ 20-min session of open- Wield for CTRL (n=8), EXP (n=12) and PSD (n=11) mice. (A) Travelled distance in open-field during 20-min session. (B) Evolution of travelled distance in open-field during the 20 minutes of the test. (C) Mean number of rearings during the 20-min session. (D) Percentage of distance travelled in the central area of the open-field. CTRL: control mice; EXP: 5G-exposed mice; PSD: Pseudo-exposed mice. Errors bars represent ± SEM. * indicates significant difference between groups of mice (p<0.05).

### Chronic 5G-exposure does not alter object / object-place recognition memory

To assess long-term memory in the OL and NOR tasks, the mice were first submitted to sessions of acquisition allowing them to explore three different objects for 15 minutes (Figure 2B, days 5 and 7). During these acquisition sessions, the CTRL, 5G, and PSD- exposed mice showed similar object exploration during the 15-min sessions (Supplementary Figure S1B-C, time to explore the three objects during the acquisition for OL: F(2,26)=0.20; P=0.81; for NOR: F(2,27)=0.03; P=0.96). Memory tests were performed 24h after acquisition by moving one of the three objects (OL task), or replacing one familiar object by a novel object (NOR task, Figure 2B). Because mice are spontaneously attracted by novelty, we analyzed the time spent exploring the displaced object for the OL task, and the novel object for the NOR task. As shown in Figure 4, 5G-exposed-mice showed preferential exploration of the displaced / novel object compared with not displaced/familiar objects (OL task: t=2.533; p=0.032; NOR task: t=3.63; P=0.0046), and their times exploring the displaced / novel object were similar to those of the control and PSD-exposed mice (OL: F(2,26)=0.1985; P=0.8212; NOR: F(2,27)=0.2080; P=0.8135).

**Figure 4.**
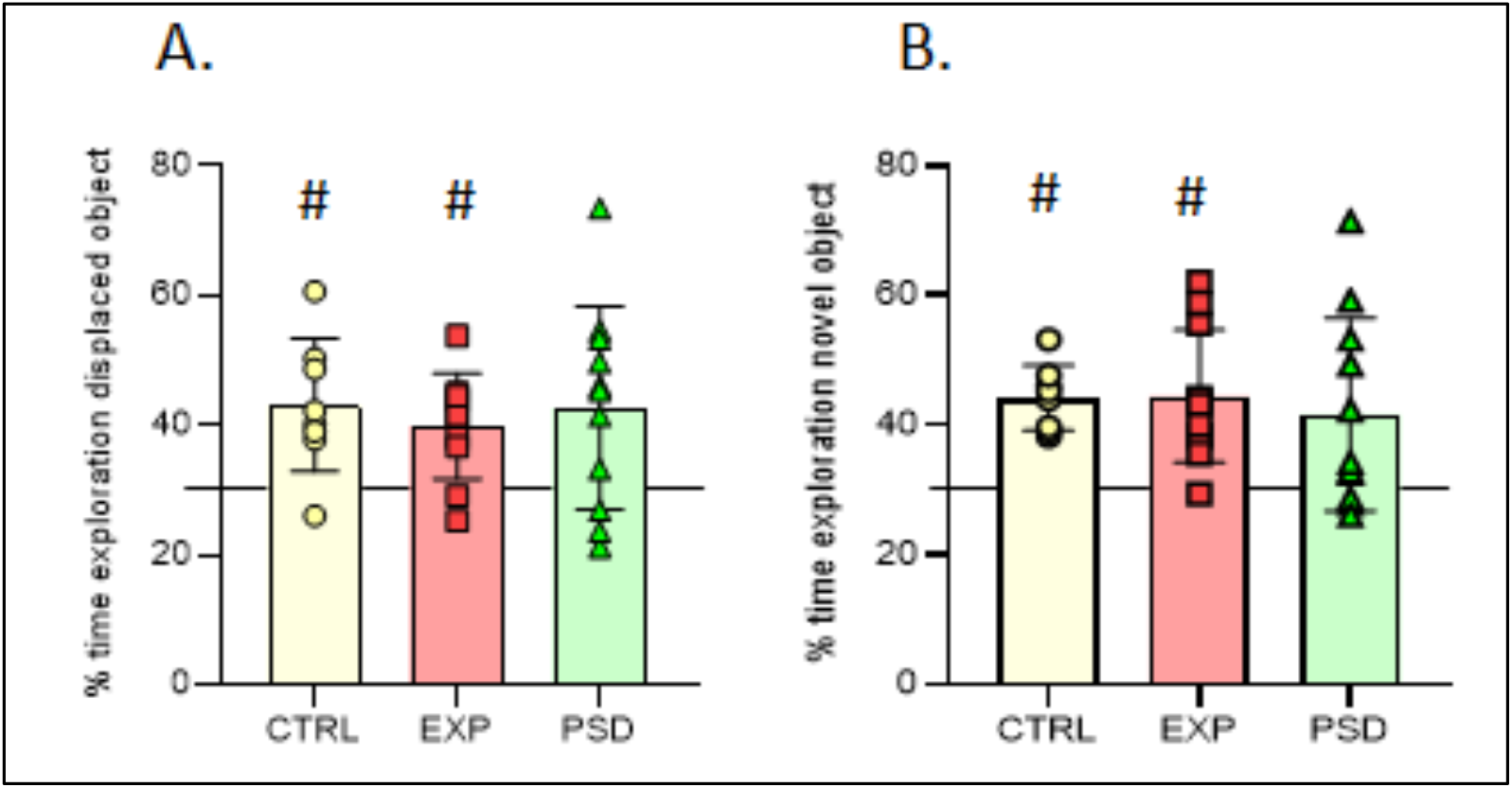
Memory tests. (A) Percentage of time spent exploring the displaced object during 5-min test session in CTRL (n=8), EXP (n=10) and PSD (n=11) mice. (B) Percentage of time spent exploring the novel object during 5-min test session in CTRL (n=8), EXP (n=11) and PSD (n=11) mice. Values are means ± SEM. The horizontal line at 33.33% indicates equal exploration of the displaced/novel and not displaced/familiar objects. **#** p<0.05 compared to 33.33% (chance). CTRL: control mice; EXP: 5G-exposed mice; PSD: Pseudo-exposed mice.

These findings demonstrate that long-term object recognition memory and long-term memory of the spatial location of objects is not affected by the 27 hours of 5G chronic exposure.

### Chronic 5G-exposure affects gene expression in the cerebral cortex

To determine the effect of the 5G-chronic exposure on gene expression in the cerebral cortex, we analyzed the coding transcriptome in two symmetrical ventral brain regions, the right and the left Ent-Pir Cx (Figure 1C3) in which the mean SAR levels were 0.43 W/kg and 0.14 W/kg, respectively. mRNA sequencing was carried out from tissue collected from 5G-exposed (n=7) and pseudo-exposed (n=8) animals that had been submitted to 27h of exposure or PSD-exposure over 6 weeks.

In PSD-exposed animals as expected, the transcriptome profiles in the right and left Ent- Pir Cx were almost identical. Significant differences in the levels of transcripts (FC >1.2, FDR-adjusted p<0.05) between the right and left Ent-Pir Cx in pseudo-exposed mice were restricted to 5 genes (Supplementary Table S2) over a total 12423 genes which expression could be detected in these brain regions.

Comparison of 5G-exposed and PSD-exposed animals revealed significant genes modulations triggered by the 5G-3.5 GHz signal. In the right Ent-Pir Cx, changes in the level of expression (FC >1.2, FDR-adjusted p<0.05) were limited to 77 genes, e.g. less than 0,7 % of the expressed genes in this CNS region. These DEG comprised 40 up-regulated and 37 down-regulated-genes in response to the 5G-chronic exposures (Figure 5A; supplementary Table S3). In the left Ent-Pir Cx, the total number of DEG was slightly higher (84 genes), despite a lower SAR level compared with the right Ent-Pir Cx. These 84 DEG segregated into 30 up-regulated genes and 54 down-regulated genes in response to the 5G exposure (Figure 5B; Supplementary Table S4). Unsupervised clustering showed clear segregation between 5G-exposed and PSD-exposed animals based on significant differences in gene expression (Figure 5 C, D). Of note, there was no effect of the 5G- exposure on the 5 genes whose expressions differed between the right and the left Ent- Pir Cx in PSD-exposed animals. Strikingly, the identity of the DEG markedly differ between the right and left Ent-Pir Cx, which only shared 8 of their DEG (Figure 5 A, B), seven of which were down regulated both in the right and the left Ent-Pir Cx.

**Figure 5.**
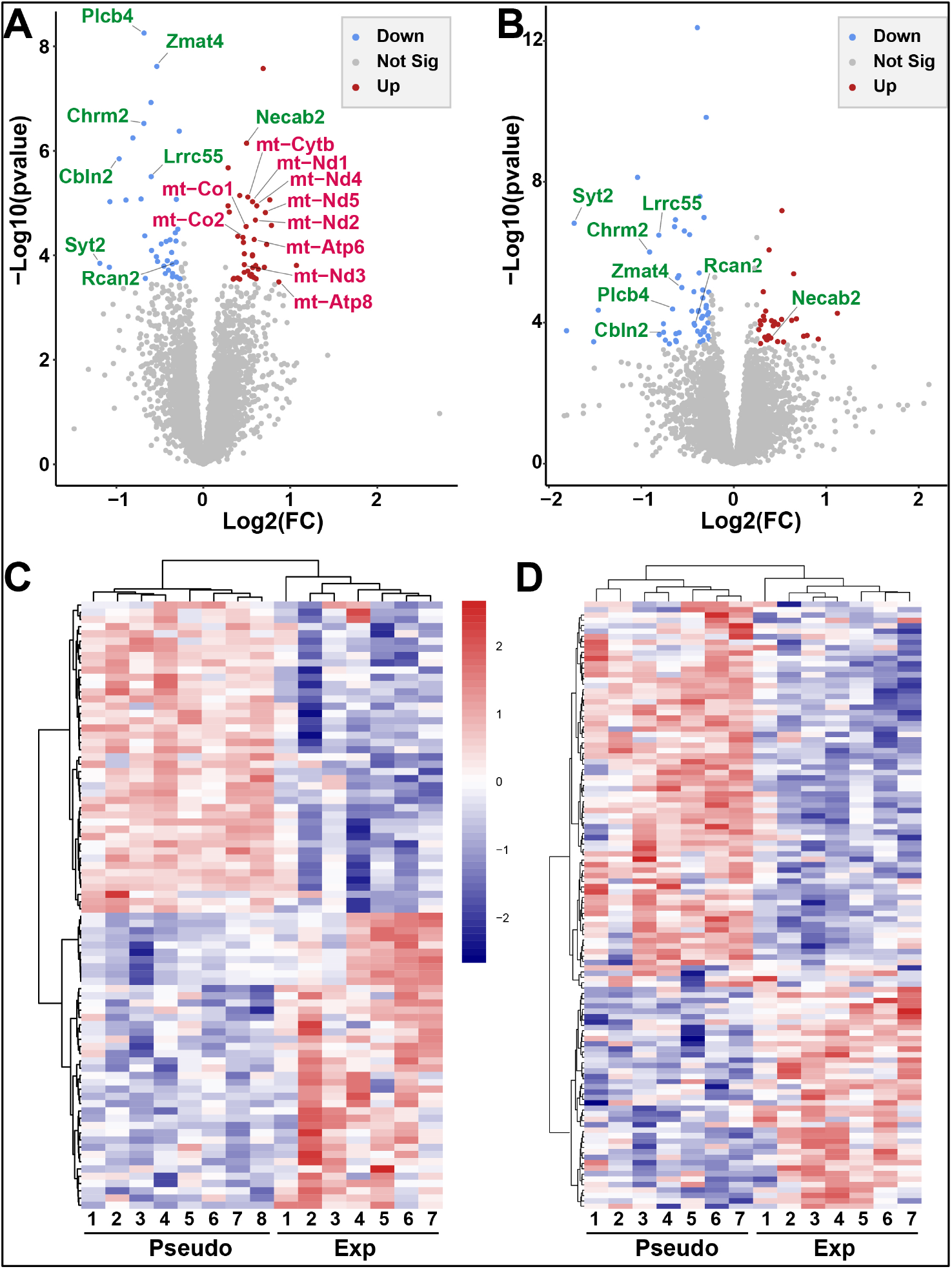
RNA-seq analysis of the Ent-Pir Cx. (A, B) Volcano plot of differential gene expression between 5G-exposed and pseudo - exposed animals (FC cutoff 1.2), in the right (A) and the left (B) Ent-Pir Cx. The genes that were significantly upregulated or downregulated in response to the 5G exposure are indicated by red or blue dots, respectively. Gray dots correspond to genes that were not significantly modulated. Upregulated gene encoded by the mitochondrial (mt) genome are written in red. Up- or downregulated genes common to the right and the left Ent-Pir Cx are written in green. (C,D) Heat maps showing color-coded normalized expression levels of the genes, which were significantly modulated by the 5G exposure (FDR adjusted p value ≤ 0.05) in the right (C) or the left (D) Ent-Pir Cx. Gene clustering was performed using Euclidean distance and ward agglomeration method. Pseudo (1–8) correspond to each of the pseudo-exposed mice; Exp(1–7) indicate 5G-exposed mice.

To specify biological processes, cellular components or molecular functions potentially affected by the 5G-triggered genes modulation, we performed a gene ontology (GO)-based enrichment analysis of the DEG.

As for the right Ent-Pir Cx, significant enrichments in GO terms were mostly related to the mitochondrial oxidative phosphorylation system (OXPHOS) [30] (Figure 6A). The enrichment in mitochondrial terms corresponded to an over-representation of genes of the mitochondrial (**mt**) genome among the DEG (Figure 5A). The mt genome contains 13 genes encoding protein subunits of the ATP-producing OXPHOS [31]. Ten of these mt-genes were upregulated (1.38 <FC< 1.83) upon exposure to the 5G signal (Figure 6B). These genes encode core subunits of enzymes complexes I, III IV and V of the OXPHOS (Supplementary Table S3), which are embedded in or anchored to the inner mitochondrial membrane [30]. Three of the mt-genes encoding peptides (mt-CO3, mt-ND4L and mt- ND6) were not identified as DEG, given our stringent criterion for statistical significance, which set the threshold for FDR-adjusted p values to 0.05. However, mt-CO3 (FC=1.35; FDR-adjusted p=0.059), mt-ND4L (FC=1.48; FDR-adjusted p=0.122) and mt-ND6 genes (FC=1.46; FDR-adjusted p= 0.142) genes showed a clear trend toward up-regulation in the right Ent-Pir Cx of 5G-exposed animals.

**Figure 6.**
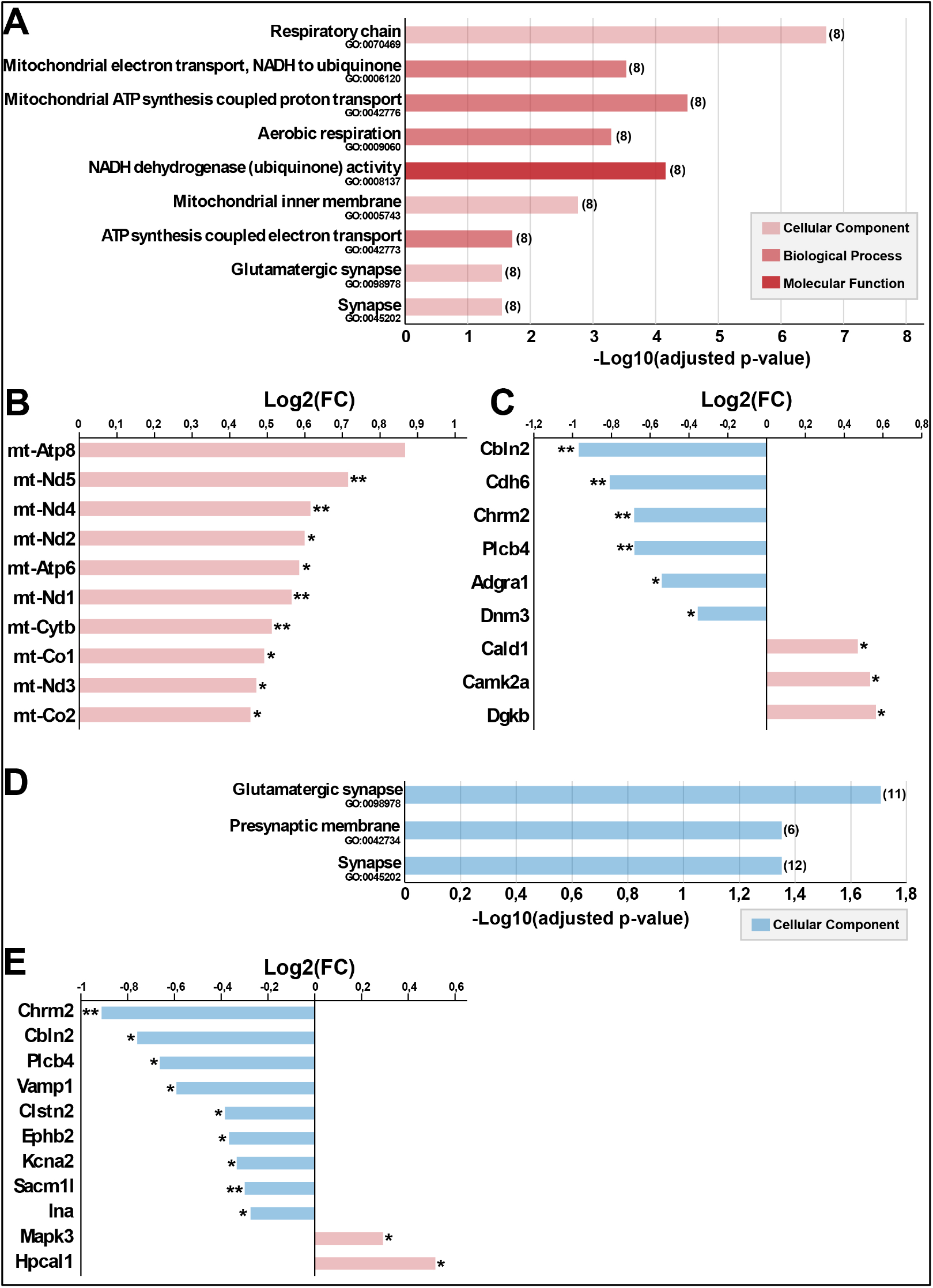
Gene ontology (GO) annotation of the differentially expressed genes (DEGs) between 5G exposed and pseudo-exposed mice. (A-C) Right Ent-Pir Cx. (A) Significantly enriched GO terms and enrichment p values are indicated. The number of DEG related to the GO terms is in brackets. (B,C) DEG associated to mitochondria-related GO terms (B) or to the “glutamatergic synapse” term (C) DEG names and fold changes (Log2( FC) are indicated. (D,E) Left Ent-Pir Cx. (D) enriched terms and p values; (E) DEG associated to the glutamatergic synapse GO term.

In both the right and the left Ent-Pir Cx, GO-based analyses highlighted significantly enriched terms defining cellular components involved in neurotransmission, such as synapses or glutamatergic synapses (Figure 6A,D). These enrichments arose predominantly from down-regulation but also from up-regulation of genes (Figure 6C,E). In the right and left Ent-Pir Cx, nine and eleven DEG encoded components of glutamatergic synapses, respectively. However, only three of these DEG (Chrm2, Plcb4 and Cbln2) were common to the right and left Ent-Pir Cx (Figure 6C,E).

Together, our results show that the chronic head exposure to the 5G -3.5 GHz signal leads to altered expression of genes in the cerebral cortex. The identity of the DEG markedly varies when comparing two homologous cortical regions that differ owing to the local SAR values during the 27h of exposure.

## Discussion

The recent launch and expansion of 5G telecommunication systems urges for assessing the brain impact of the EMF generated with this technology. We have set up an asymmetrical head exposure system to a calibrated 5G modulated-3.5 GHz signal, which was adapted to awake adult mice. Dosimetric analyses indicate that the brain average SAR level was 0.19 W/kg whereas maximal SAR levels in the range of 0.5 W/kg were reached in the right Ent-Pir Cx. We have investigated the effect of repeated head exposures on the mouse CNS through assessment of animal behaviors and transcriptome profiling. Our results show no significant effect of the 5G signal on locomotion, anxiety level and memory capacities, but they reveal biological responses quantified by modulations of gene expression in the cerebral cortex.

At the behavioral level, we have analyzed the influence of a 5G-3.5 GHz signal on mice, assessing the locomotion and exploration in an OF. None of the analyzed behavioral parameters such as travelled distance, numbers of rearings or distance in the central area revealed any significant alteration in the exploration and anxiety level, when applying the 5G signal 1h daily, 5 days a week over a 6-week period. Similarly, we found no significant effect of the 5G signal on memory abilities as determined in object-location and novel-object recognition tasks. Our results differ from those of a recent study showing that 1h daily whole body exposures of adult mice to pulsed 3.5 GHz RF carried out over 35 days led to increased anxiety levels, albeit without modification of learning or memory performances [32]. However, this report did not specify the pulse mode that was applied to the exposures nor the SAR levels that were reached in the brain or the body of the exposed mice. So far, to our knowledge, no other rodent study has investigated how a chronic exposure to 3.5 GHz RF during adulthood might affect emotional states or memory capacities. However, increases in anxiety levels were detected in adult zebrafishes that were exposed to 3.5 GHz for 42h at a high SAR level (8.7 W/kg) during embryonic stages of life, as indicated by enhanced escape response to predators [15]. Zebrafish embryos exposed for 1 or 4h to 3.5 GHz RF at an estimated SAR of 1.1 W/kg subsequently displayed anxiety-like behavior during larval stage of life, which was marked by modifications in the thigmotaxis or wall-hugging behavior triggered by visual or auditory stimuli [16]. Behavioral impacts of RF-EMF depend on various parameters including not only the number, the duration and the power of the exposures but also the type of modulation applied to the emitted RF which changes according to the generation of mobile communication systems [3,33,34]. Our study provides a first analysis of the effect of a 5G-modulated 3.5 GHz signal on anxiety or memory capacities in mammals.

Transcriptome profiling, was previously used to determine gene responses in the brain of rodents that were exposed to GSM-related EMF [35–40]. Here we found that the chronic 5G-3.5 GHz exposure, triggered significant modulations in less than 1% of the expressed genes in the right Ent-Pir Cx, where the local SAR level reached its maximal levels.

Expectedly, in the absence of EMF exposure, the right and left Ent-Pir Cx displayed virtually identical transcriptome profiles (differences in gene expression limited to 5 over more than 12 000 expressed genes in pseudo-exposed animals). Due to the asymmetrical head exposure, we compared the gene responses in the right and left Ent-Pir Cx, which were differentially exposed to the 5G-3.5 GHz signal. We observed that a 3-time decrease of the mean SAR level was associated to marked changes in the identity of the DEG without reduction in the number of responsive genes. In fact, the number of DEG was higher in the left Ent-Pir Cx (84 DEG, mean local SAR of 0.14 W/kg) than in the right Ent-Pir Cx (77 DEG, mean SAR: 0.43 W/kg), whereas only 8 DEG were shared by the right and left cortical areas. This result indicates that the magnitude of the gene response does not continuously increase with the level of energy absorbed in the tissue, when SAR values range between 0.1 and 0.5 W/kg. EEG studies performed with human subjects indicate that visual activities impact on the cortical cell responses triggered by a low power-head exposure to a 5G-3.5 GHz signal [8]. We speculate that differences in SAR values and neuronal activities concurred in shaping different gene responses in the right and left Ent-Pir Cx.

Despite the marked differences in gene identities, the DEG profiles in the right and left Ent-Pir Cx shared significant enrichments in genes related to glutamatergic synapses that convey defined excitatory neurotransmissions in these cortical areas [41]. The entorhinal cortex is the primary spatial input to the hippocampus and bridge neuronal signaling between hippocampus and cortical regions including the piriform cortex. Neuronal activity in the entorhinal cortex plays major role in spatial memory and navigation (reviewed in [42,43]) and was also proposed to be involved in object recognition memory [44]. Our current behavioral analyses show no alteration of these memory capacities in the animals exposed to the 5G-3.5GHz signal. However, this does not discard the possibility that the synapse-related DEG responses reflected alterations in excitatory neurotransmission, which remained too weak to affect memory performances. In line with this hypothesis, altered neuronal activity sparing cognition was recently documented in a study combining behaviors assessment and intracerebral cFos imaging in rats submitted to subchronical exposure to a GSM-900 MHz signal (45 min exposure daily over two weeks at a brain averaged SAR of 1 or 3.5 W/kg). The GSM- exposures led to reduced neuronal activity in cortical and hippocampal areas involved in memory tasks without modification in memory performances [45]. Furthermore, suppressive effects of RF-EMF on glutamatergic neurotransmission or neuronal bursting activity were observed in culture studies in which rat cortical or hippocampal neurons were exposed to GSM-1800 MHz microwaves at SAR levels ≥ 0.1 W/kg [46,47], as well as in vivo, in rats submitted to acute head-exposures to GSM-900 or 1800 MHz or LTE-1800 MHz signal [17,20,48,49].

While our study highlights synapse-related DEG in both the right and left ent-Pir Cx , we found that the DEG profile in the right Ent-Pir Cx differed from the left counterpart by an over representation mt genes (supplementary Table S3, figure 6B). The mt genome is a double strand circular DNA with cellular copy numbers that vary between few hundred and several thousand according to cell types. It contains 37 intronless genes that encode 13 peptides, 22 transfer RNAs and 2 ribosomal RNAs involved in the translation of these peptides. Mitochondrial transcription generates polycistronic transcripts from which the mature mRNA, transfer RNAs and ribosomal RNAs are derived, resulting in jointed transcription of different mt genes [31,50]. Each of the 13 mt-genes encoding peptides displayed significant or clear trends toward upregulation in the right Ent-Pir Cx of 5G- exposed mice. This suggests that the 5G exposures resulted in transcription-sparing duplication or enhanced transcription of the mt-DNA. The mitochondrial genome encodes essential components of the OXPHOS, as illustrated by pathological consequences of mt-gene mutations [51,52]. The OXPHOS comprises five protein complexes, four of which (complexes I to IV) ensure a stepwise series of transfer of electrons from reducing components to molecular oxygen. The process generates a transmembrane electrochemical gradient, which is the driving force for several mitochondrial functions including the synthesis of ATP carried out by complex V (FoF1 ATP synthase) [30,53]. Reactive oxygen species (ROS) are common side products of oxidative phosphorylation, which predominantly originate from the activity of complexes I and III and can be source of oxidative damage to DNA, proteins and lipids [54,55].

Numerous studies have investigated the capacity of RF-EMF to promote ROS generation and tissue oxidative stress (reviewed in [56]). Increased ROS activity has been observed in the brain of rodents chronically exposed to RF-EMF at different frequencies (900, 1800, 2100, and 2450 MHz) used for 2-4G or WIFI communications, applying whole-body SAR values in the range from 0.6 10^-3^ to 0.9 W/kg [57–62]. In some studies, EMF-induced oxidative stress was associated with impaired spatial memory [62] or DNA damage [58,59,61].

With respect to the 3.5 GHz frequency band, Bektas et al. (2022) [63] found that repeated head exposures of adult rats (2h/day for a 1month) to a GSM-modulated 3.5 GHz resulted in enhanced levels of brain ROS that were assessed in whole brain homogenates, whereas SAR values were estimated to 323 mW/kg in the gray matter of the brain. Yang et al. (2022) [11] observed oxidative brain lesions marked by lipid peroxidation and swollen mitochondria, when applying 3.5 GHz continuous wave to adult guinea pigs for 3 days, with whole body SAR values ≥ 2W/kg. Increased ROS productions were also observed in cultured human astrocyte or glioma cell lines that were exposed to high power 3.5 GHz pulses [64].

In line with the hypothesis that a 5G modulated 3.5 GHz signal could promote ROS production, our study shows that the exposure to the 5G-3.5 GHz signal increases expression of mt-genes encoding sub units of four of the five OXPHOS protein complexes (supplementary Table S2) including the ROS-producing complexes I and III. However, the OXPHOS proteins complexes involve the assembly of around 90 protein subunits, which have a dual mitochondrial and nuclear origin. Most of the subunits are encoded by nuclear genes and imported into the mitochondria [30]. Our RNA seq analyses show no significant change in the expression of these nuclear genes. Furthermore, the cellular distribution of the mt-gene response to the 5G-3.5GHz signal is not known. This issue may have functional relevance, considering that the molecular assembling of the OXPHOS complexes differ between neuronal and astroglial mitochondria, such that the production of ATP is more efficient in neurons whereas ROS production is more prominent in astrocyte [65]. It remains to determine whether the upregulation of mt-genes observed here reflects metabolism adaptations marked by enhanced production of ATP or an oxidative stress, which could affect the right Ent-Pir Cx in the animal exposed to the 5G-3.5 GHz signal.

## Conclusion and limitations

Altogether, our results show that daily 1h head exposure to a 5G-3.5 GHz over a 6 weeks period, does not alter emotional state and memory performances, but trigger significant modifications of expression in a limited set of genes, which can potentially affect glutamatergic synapses and mitochondrial activities. We acknowledge limitations to our study. We cannot exclude that prolongations of the head exposures beyond 6 weeks could ultimately affect the emotional state or memory abilities of the exposed mice. The experiments were carried out with male mice, because of the impossibility to house separately male and female mice during 5G exposures and behavioral testing. Further investigation will be required to evaluate whether female and male mice could be differentially affected by chronic exposure to a 5G-3.5 GHz. In addition, our RNA seq analyses was performed at a single time point e.g. 24h after the last exposure to the 5G-3.5 GHz signal. The kinetic and the reversibility of the reported changes in gene expression is undetermined. The 5G-triggered transcriptome modifications were observed in cortical areas where the average SAR levels range around 0.43 or 0.14 W/kg. These SAR levels may be considered in light of the European safety guidelines for human head exposures [66], which set the upper SAR limit to 2 W/kg. This value is higher than the corresponding SAR levels reached in the brain due to the energy absorption in the surrounding skull tissues. Recent dosimetric analyses of human exposures to downlink RF-EMF from base stations shows intracortical SAR levels attributed to environmental 5G-3.5 GHz that are much lower than values applied in our study, being less than 1 mW/kg [9]. Further investigations are needed to specify levels of SAR reached in the human cerebral cortex when mobile phones emitting a 5G-3.5 GHz signal are hold close to the ear of the mobile phone user.

## Fundings

This research was supported by the French National Research Program for Environmental and Occupational health of ANSES (grant 2020/2 RF/14).

## Institutional Review Board Statement

The experiments were performed under the national license A-91-557 (APAFIS project # 33606 from the “Direction Générale de la recherche et de l’innovation”) using procedure No. 32-2011 validated by the Ethics Committee in Animal Experimentation (CEEA 59, Paris Centre et Sud). All procedures were performed in accordance with the guidelines established by the European Communities Council Directive (2010/63/EU Council Directive Decree).

## Availability of data and materials

The RNA seq datasets generated and analyzed during the current study will be available in the GEO repository at the time of publication

## Acknowledgements

We thank the Paris Brain Institute’s Data Analysis Core for assistance in the bioinformatic analyses

## Conflicts of Interests

The authors have no conflicts of interests to declare

---

**Supplementary Figure S1.**
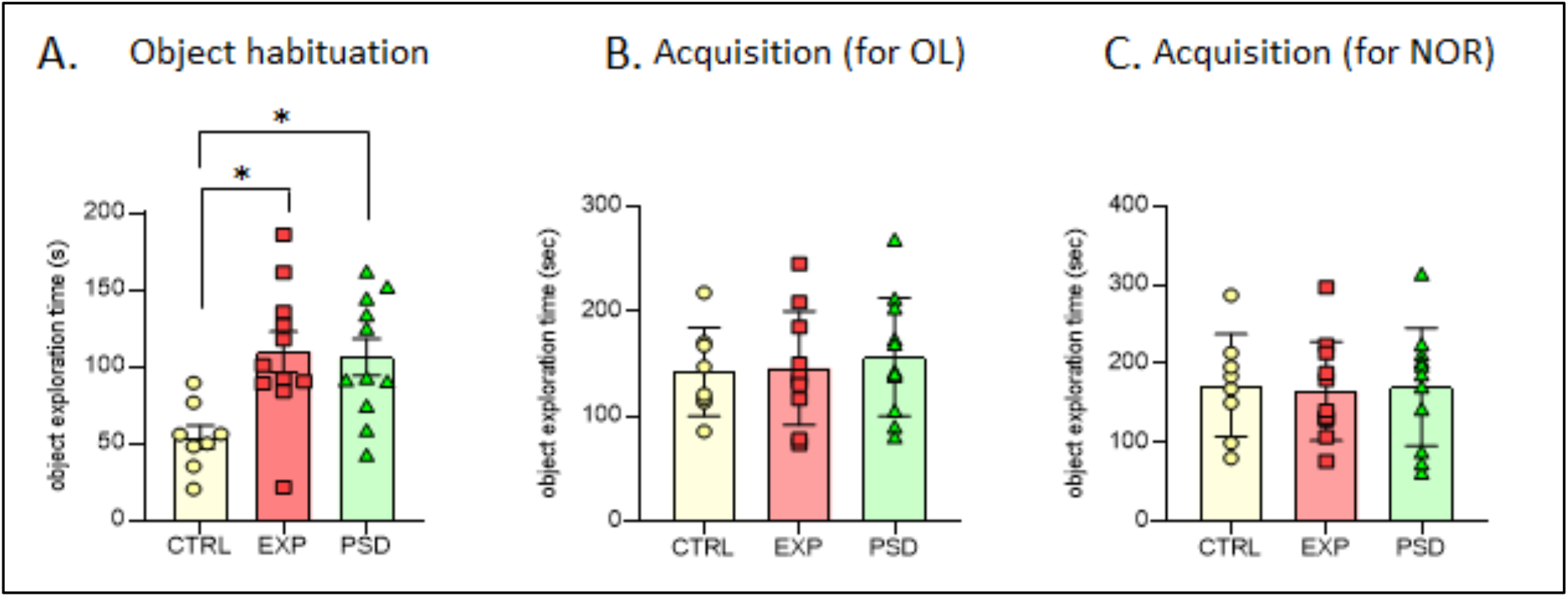
(A) Object time exploration during the object habituation session (day 4) for CTRL (n=8), EXP (n=11) and PSD (n=11) mice. (B-C) Object time exploration during the acquisition sessions (days 5 or 7) for CTRL (n=8), EXP (n=10/11) and PSD (n=11) mice. CTRL: control mice; EXP: 5G-exposed mice; PSD: Pseudo-exposed mice. * indicates significant difference.

**Supplementary Table S1:** Behavioral parameters observed during the 2^nd^ and 3^rd^ session in openfield (OF) and statistical results (factorial ANOVA). CTRL: control mice; EXP: 5G- exposed mice; PSD: Pseudo-exposed mice.

|  | CTRL | EXP | PSD | ANOVA |
| --- | --- | --- | --- | --- |
| **2<sup>nd</sup> session - OF** |  |  |  |  |
| Travelled distance | 57.14 ± 3.47 | 57.75 ± 1.98 | 54.81 ± 2.76 | F(2,28)=0.36; P=0.69 |
| Numbers of rearings | 79.13 ± 7.06 | 93.83 ± 5.99 | 88.09 ± 6.72 | F(2,28)=1.16; P=0.32 |
| Travelled distance in center (%) | 46.15 ± 2.28 | 50.17 ± 2.08 | 50.11 ± 2.3 | F(2,28)=0.91; P=0.41 |
| **3<sup>rd</sup> session - OF** |  |  |  |  |
| Travelled distance | 52.13 ± 1.9 | 54.74 ± 2.43 | 58.4 ± 3.27 | F(2,28)=1.22; P=0.30 |
| Numbers of rearings | 94.5 ± 7.91 | 97.42 ± 5.21 | 100.2 ± 6.39 | F(2,28)=0.18; P=0.83 |
| Travelled distance in center (%) | 45.14 ± 3 | 43.44 ± 1.65 | 50.28 ± 2.82 | F(2,28)=2.26; P=0.12 |

**Supplementary Table S2.** - Genes Differentially Expressed in the Right and the Left Ent-Pir CX in Pseudo-exposed Animals.

| Gene symbol | Gene Name | Expression Right versus Left Ent-Pir CX | Fold-change | adjusted p value |
| --- | --- | --- | --- | --- |
| Ankfn1 | ankyrin-repeat and fibronectin type III domain containing 1 | Down | 2,001998642 | 0,042075965 |
| Gnb4 | guanine nucleotide binding protein (G protein), beta 4 | Down | 1,44379541 | 0,042075965 |
| Igfbp6 | insulin-like growth factor binding protein 6 | Down | 1,703388817 | 0,042075965 |
| Ptgs2 | prostaglandin-endoperoxide synthase 2 | Down | 1,849700874 | 0,042075965 |
| Smoc2 | SPARC related modular calcium binding 2 | Down | 1,717193717 | 0,042075965 |

**Supplementary Table S 3.**
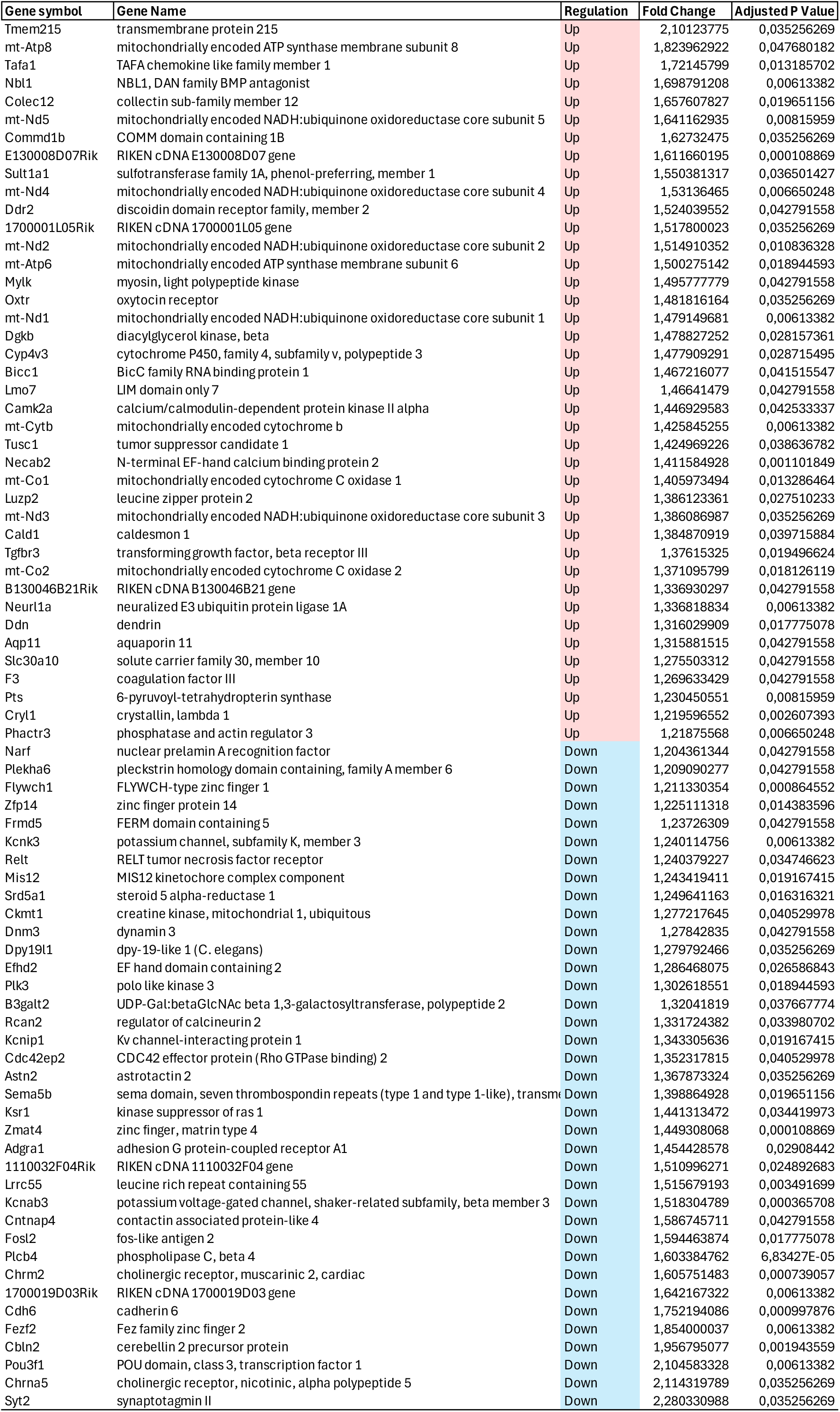
- Genes Differentially Expressed in the Right Ent- Pir CX of 5G/3.5GHz-exposed versus Pseudo-exposed Mice.

**Supplementary Table S4.**
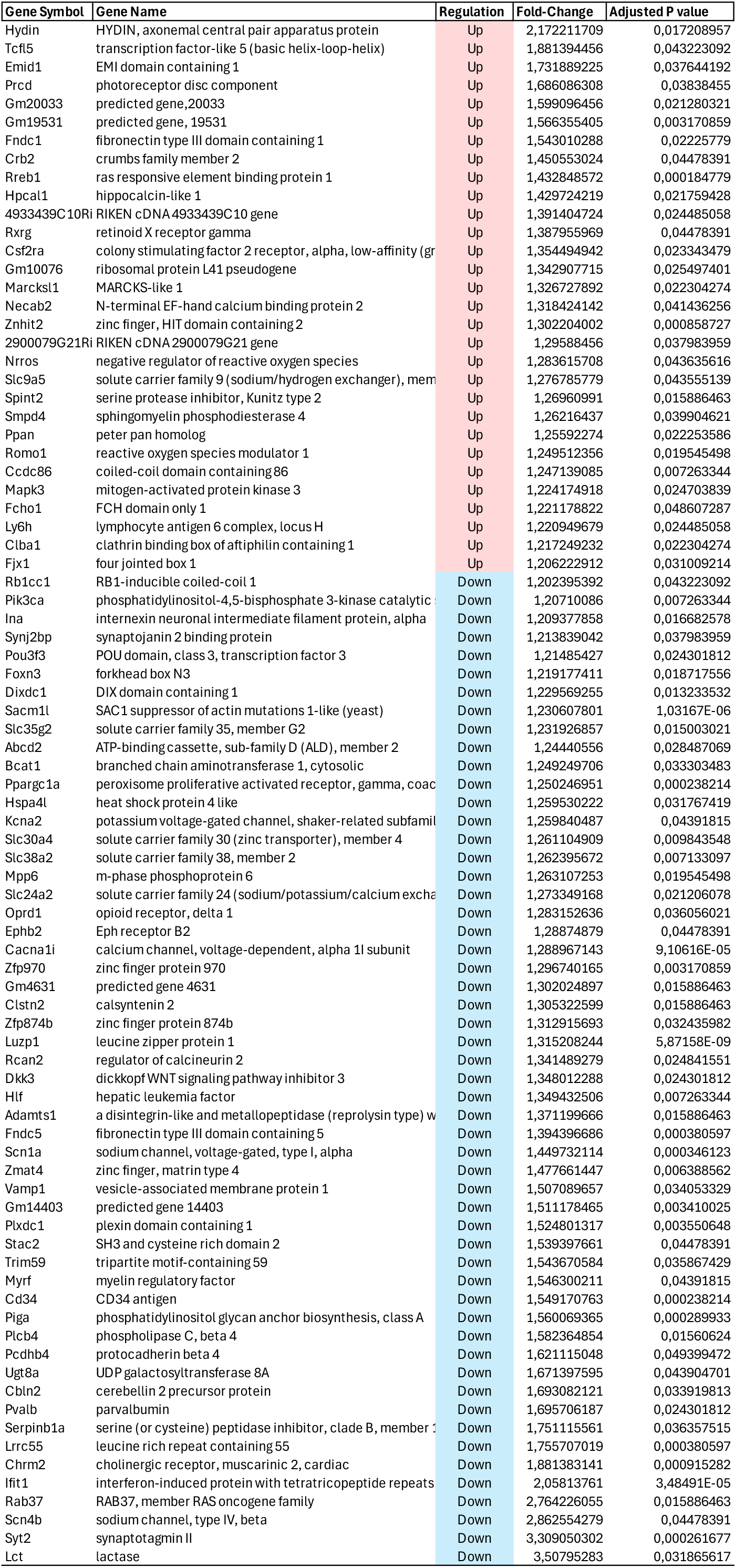
- Genes Differentially Expressed in the Left Ent- Pir CX of 5G/3.5GHz- exposed versus Pseudo-exposed Mice.

## Notes

### Competing Interest Statement

The authors have declared no competing interest.

